# C3D: A tool to predict 3D genomic interactions between cis-regulatory elements

**DOI:** 10.1101/197301

**Authors:** Tahmid Mehdi, Swneke D. Bailey, Paul Guilhamon, Mathieu Lupien

## Abstract

**Motivation:** The 3D genome architecture influences the regulation of genes by facilitating chromatin interactions between distal cis-regulatory elements and gene promoters. We implement Cross Cell-type Correlation based on DNA accessibility (C3D), a highly customizable computational tool that predicts chromatin interactions using an unsupervised algorithm that utilizes correlations in chromatin measurements, such as DNaseI hypersensitivity signals.

**Results:** C3D accurately predicts 32.7%, 18.3% and 24.1% of interactions, validated by ChIA-PET assays, between promoters and distal regions that overlie DNaseI hypersensitive sites in K562, MCF-7 and GM12878 cells, respectively.

**Availability:** Source code is open-source and freely available on GitHub (https://github.com/LupienLabOrganization/C3D) under the GNU GPLv3 license. C3D is implemented in Bash and R; it runs on any platform with Bash (≥4.0), R (≥3.1.1) and BEDTools (≥2.19.0). It requires the following R packages: GenomicRanges, Sushi, data.table, preprocessCore and dynamicTreeCut.

## INTRODUCTION

Genome-wide association studies (GWAS) have highlighted that non-coding variants play a large role in the etiology of common diseases by targeting cis-regulatory elements (CREs), mainly enhancers as opposed to promoters (Cowper-Sal·lari et al., 2012; Maurano et al., 2012; Schaub et al., 2012). However, it can be difficult to identify the target genes of these CREs. The challenge arises because enhancer-type CREs are often distal to their target genes relying on chromatin interactions to come in physical proximity with promoters, often skipping over the most proximal gene (Dekker et al., 2013; Sanyal et al., 2012). Means to delineate the 3D architecture of the genome are therefore required.

There are several assays for detecting chromatin interactions based on Chromosome Conformation Capture (3C) (Dekker et al., 2002), such as Chromatin Interaction Analysis with Paired-End-Tag sequencing (ChIA-PET) (Fullwood et al., 2009), Hi-C, 4C and 5C (Göndör et al., 2008; Dostie et al., 2006; van Berkum et al., 2010). These methods require large amounts of material, not available for each cell or tissue of interest, and can be limited in resolution. Computational approaches to predict chromatin interactions present an attractive alternative or complementary approach. Ernst et al. (2011) and Corradin et al. (2013) proposed methods that utilize Pearson correlation coefficients between chromatin modification levels at CREs across a collection of cell types. Sheffield et al. (2013) applied a similar correlation method but comparing chromatin accessibility signals with gene expression levels. Thurman et al. (2012) predicted interactions from correlations based solely on chromatin accessibility across DNaseI hypersensitive sites (DHSs) using aggregated DNaseI hypersensitivity signals across 79 ENCODE cell lines (ENCODE Project Consortium, 2012). This allowed the pairing of distal CREs with active promoters as opposed to genes, which facilitated direct comparison with in vivo 3C-based assays. This method reported a significant enrichment of chromatin interactions called by ChIA-PET (p < 10^-15^) and 5C (p < 10^-13^) in K562 cells with highly correlated (r > 0.7) promoter-distal predicted interactions. Here, we expanded this approach with our Cross Cell-type Correlation in DNA accessibility (C3D) tool to predict chromatin interactions between CREs by expanding the analysis to an additional 100 biological samples [47 cell lines from Duke University (ENCODE Project Consortium, 2012) and 53 cell-types from the University of Washington (UW) (Roadmap Epigenomics Consortium et al., 2015)] into its correlation calculations and by applying statistical tests to identify significant correlations.

## METHODS

C3D computes correlations between genomic measurements to predict chromatin interactions between genomic regions of interest. For instance, these genomic measurements can be DNase-seq (Boyle et al., 2008) or ATAC-seq (Buenrostro et al., 2013) signal intensities. It requires a reference catalogue of genomic coordinates found in a cell or tissue of interest (ie. all regions of accessible chromatin in a given cell or tissue type). It also requires an anchor file containing genomic coordinates of interest such as the promoters of genes of interest. Lastly, it requires signal files from a set of biological samples; these files should include genomic coordinates and their corresponding measurements.

As a case example, we used DNase-seq from ENCODE and the Roadmap Epigenomics Project (ENCODE Project Consortium, 2012; Roadmap Epigenomics Consortium et al., 2015) for the signal files. For each DNase-seq signal file, the maximal DNaseI signals are computed at each region in the reference catalogue. Quantile normalization (Bolstad et al., 2003) is then applied to ensure identical scaling of signals across signal files. The DynamicTreeCut algorithm (Langfelder et al., 2008) is then used to cluster the signal files based on the similarity between their signals across the reference catalogue. Each region in the reference catalogue receives a vector of consensus genomic measurements where each measurement corresponds to a cluster. The consensus genomic measurement of a cluster is the average signal from its respective signal files. This limits biases introduced by an overrepresentation of a particular tissue/cell-type. C3D then iterates through the sites listed in the reference catalogue, which overlap a region from the anchor file. For each of these sites, the Pearson correlations between the consensus genomic measurements of each reference catalogue genomic coordinates and those in the anchor file within a user-defined window are computed. For each pair of sites, a two-sided t-test is performed to determine whether they are significantly correlated and the p-values are adjusted through the Benjamini-Hochberg procedure (Benjamini and Hochberg, 1995). Instructions for installing and running C3D are provided in the supplementary information along with a tutorial.

We validated the predictive accuracy of C3D using RNA polymerase II (POL2) ChIA-PET datasets for K562, MCF-7 (ENCODE Project Consortium, 2012) and GM12878 (Tang et al., 2015) cells. ChIA-PET called chromatin interactions are reported as pairs of genomic regions (or paired-end-tags) mapped to GRCh37/hg19. Interactions in these datasets are filtered to focus on interactions between promoters and distal sites that were 10 to 500 kilobase pairs (kbp) apart and overlapped DHSs reported in the matched cell-type (ENCODE Project Consortium, 2012; Roadmap Epigenomics Consortium et al., 2015). We define promoters as tags that are at most 1 kbp upstream from a transcription start site defined in GENCODEv19 (Harrow et al., 2012); all other tags are considered distal. Shuffling the distances between tags and rearranging them accordingly generated a matched control set of paired-end-tags (matched Control-PETs), for each interaction dataset, to test C3D’s ability to control for false positives. We calculated correlations between promoter and distal DHSs in each of the three cell-types. For comparison, correlations were computed across DNase-seq databases of 79 cell lines from UW, 47 cell lines from Duke University (ENCODE Project Consortium, 2012), 53 cell-types from the Roadmap Epigenomics Project [also from UW] (Roadmap Epigenomics Consortium et al., 2015) and a combined database consisting of all 179 samples.

## RESULTS

In K562, MCF-7 and GM12878 cells, interacting DHSs tend to have significantly higher correlations, across ENCODE and Roadmap Epigenomics DNase-seq data, than DHS-pairs in the matched Control-PETs (Figure 1A). Correlation thresholds for each cell are chosen as the value that 5% of matched Control-PETs have correlations above. With a cut-off of 0.62 (p < 5.74x10^-7^, q < 1.68x10^-5^) C3D predicted 32.7% and 18.3% of the ChIA-PET connections in K562 and MCF-7, respectively. In GM12878, a cut-off of 0.63 (p < 3.32x10^-7^, q < 1.10x10^-6^) allowed it to have a true positive rate (TPR) of 24.1% (Figure 1A).

**Fig. 1.**
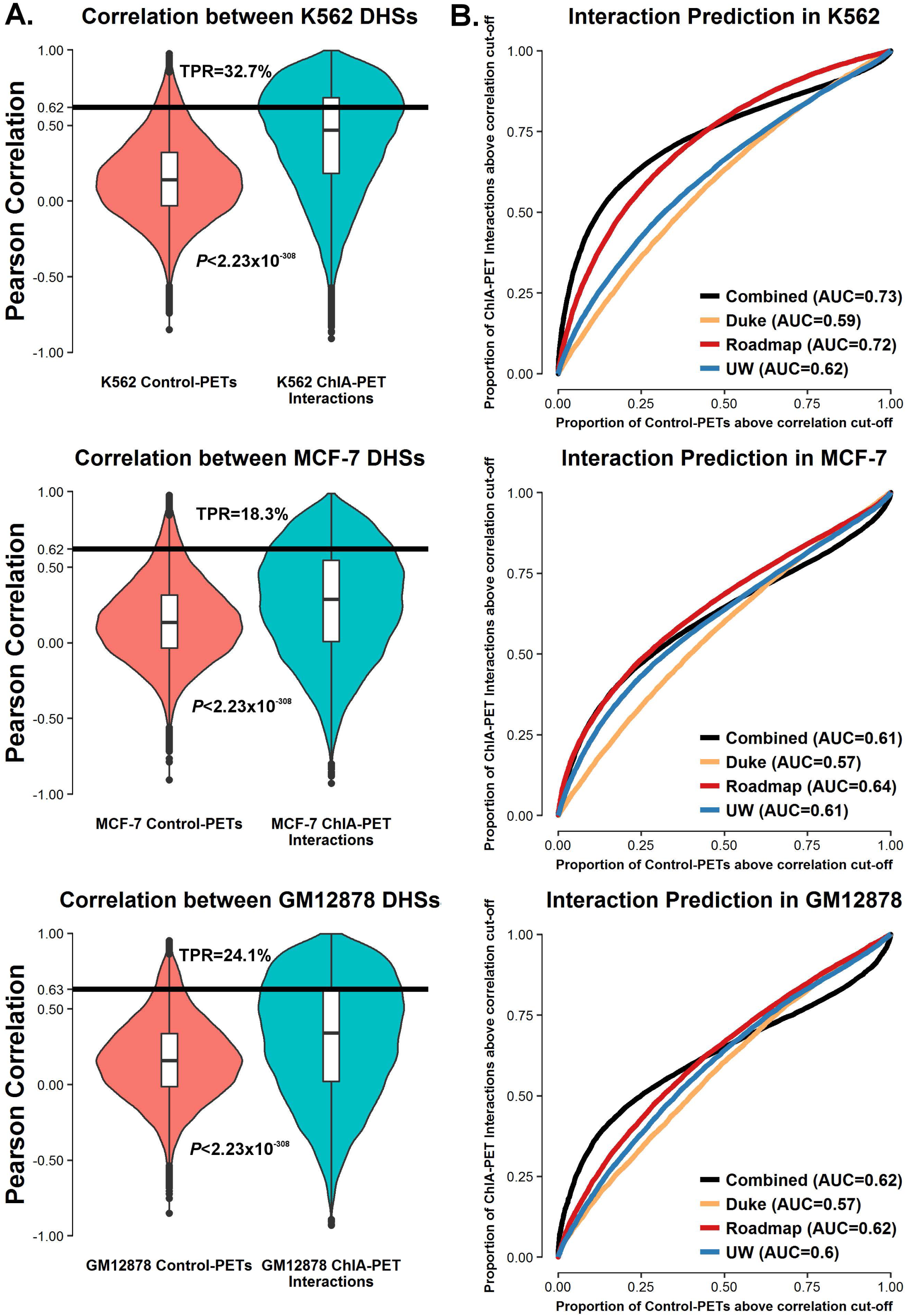
Performance of C3D on ChIA-PET and control datasets. **(A)** For the ChIA-PET data, C3D is applied to catalogues of K562, MCF-7 and GM12878 DHSs individually and calculates correlations across the combined panel of DNase-seq signals. Since some promoters and distal tags contain multiple DHSs, each paired-end tag was assigned the maximal correlation between a DHS in its promoter tag and one in its distal tag. For the control sets, C3D is applied to a merged catalogue of DHSs consisting of all DNase-seq peaks from the samples in the database. Each Control-PET received the correlation between a DHS in its promoter tag and a random one in its distal tag. For each cell-type, a Mann-Whitney U test compared the correlations of its interacting regions and control-PETs. TPRs are calculated at the correlation cut-offs that 5% of control-PETs are above. **(B)** C3D is run on the catalogues of K562, MCF-7 and GM12878 DHSs individually using four different databases: 79 samples from UW, 47 samples from Duke University, 53 samples from the Roadmap Epigenomics Project and a combined database of all 179 samples. For each of the three cell-types, the proportion of control-PETs and ChIA-PET connections classified as interactions are plotted at different correlation thresholds ranging from -1 to 1. AUCs are computed.

To compare the influences of the four aforementioned databases on the predictive abilities of C3D, receiver operating characteristic (ROC) curves were generated and areas under the curves (AUCs) are measured (Figure 1B). Calculating correlations across the combined DNase-seq database was more effective than the individual datasets in predicting interactions in K562 as it generated the highest AUC. It produced the same AUC as the Roadmap Epigenomics Project database for GM12878 but it had a greater Matthews Correlation Coefficient (MCC) (Matthews, 1975) at high thresholds (r > 0.63) (Supplementary Table 1). In MCF-7, the Roadmap Epigenomics Project database had the highest AUC but its predictive accuracy was similar to that of the combined database at high thresholds (Figure 1B). With a threshold of 0.63, C3D achieved higher MCCs in all three cells with the combined database than the individual datasets (Supplementary Table 1). Thus, C3D can predict chromatin interactions and control for false positives if adequate databases and correlation thresholds are applied.

While we present a method to predict chromatin interactions, this method will also report correlations dependent on other biological processes, such as gene co-expression across cell-types reflected by the signals at DHSs within their promoters, stressing the need for their functional validation.

## FUNDING

This work was supported by Prostate Cancer Canada and is proudly funded by the Movember Foundation (grant #RS2014-04 to M.L.), with the additional support of the Canadian Cancer Society and the Princess Margaret Cancer Foundation (M.L.). M.L holds an Investigator Award from the Ontario Institute for Cancer Research; a Canadian Institutes of Health Research (CIHR) New Investigator Award; and a Movember Rising Star Award from Prostate Cancer Canada. P.G is supported by a CIHR Fellowship (MFE 338954). S.D.B. is a Knudson and CIHR postdoctoral fellowship recipient.

## ACKNOWLEDGEMENTS

We thank Dr. Michael M. Hoffman for helpful comments and suggestions on the manuscript.

## Conflict of Interest

none declared.

